# Non-assortative community structure in resting and task-evoked functional brain networks

**DOI:** 10.1101/355016

**Authors:** Richard F. Betzel, Maxwell A. Bertolero, Danielle S. Bassett

## Abstract

Brain networks exhibit community structure that reconfigures during cognitively demanding tasks. Extant work has emphasized a single class of communities: those that are assortative, or internally dense and externally sparse. Other classes that may play key functional roles in brain function have largely been ignored, leading to an impoverished view in the best case and a mischaracterization in the worst case. Here, we leverage weighted stochastic blockmodeling, a community detection method capable of detecting diverse classes of communities, to study the community structure of functional brain networks while subjects either rest or perform cognitively demanding tasks. We find evidence that the resting brain is largely assortative, although higher order association areas exhibit non-assortative organization, forming cores and peripheries. Surprisingly, this assortative structure breaks down during tasks and is supplanted by core, periphery, and disassortative communities. Using measures derived from the community structure, we show that it is possible to classify an individual’s task state with an accuracy that is well above average. Finally, we show that inter-individual differences in the composition of assortative and non-assortative communities is correlated with subject performance on in-scanner cognitive tasks. These findings offer a new perspective on the community organization of functional brain networks and its relation to cognition.

## INTRODUCTION

The human brain is a complex network of functionally and structurally interconnected brain areas. This network exhibits non-random topological attributes that exist along a spectrum, ranging from local properties of individual brain areas to global properties reflecting the organization of the entire brain [1, 2]. Situated between these two extremes is a rich meso-scale comprising sub-networks of topologically-related brain areas called “communities” or “modules” [3, 4]. The brain’s community structure highlights patterns and regularities in its wiring diagram, delineating groups of brain areas with similar functional connectivity (FC) profiles and (presumably) shared functionality [5–7].

In the absence of explicit task instructions, the brain’s community structure reflects its intrinsic and baseline functional architecture [5, 8]. Building on this observation, a growing number of studies have begun characterizing the principles by which communities reconfigure as subjects perform cognitively demanding tasks. These task-evoked changes appear subtle, and yet also display a high degree of specifity in terms of which communities reconfigure and the tasks that prompt the reconfiguration itself [9]. Accompanying these changes are increases in network-wide integration, with inter-community FC becoming stronger as the neural activity of intrinsic brain systems required for task performance becomes correlated [10]. Importantly, the magnitude of reconfiguration or the extent to which subjects reconfigure into favorable network states has been associated with intersubject performance on cognitive tasks [11–16].

Though informative, these studies suffer from a common limitation. Namely, they all assume that the brain’s community structure is uniformly assortative; that is, communities are internally dense and externally sparse in terms of their FC. It is true that assortative communities confer evolutionary and functional advantages to nervous systems, such as specialization of function [17], evolutionary adaptability [18], and robustness to perturbations [19]. Yet, their ubiquity throughout the field of network neuroscience may also be at least partly attributable to methodological convenience. Community detection algorithms such as modularity maximization [20] and Infomap [21] can be implemented without much user input and (under certain conditions) offer fast and accurate estimates of a network’s community structure. However, both methods are built around objective functions that, when optimized, seek partitions of the network into assortative communities. Given the widespread use of these methods in network neuroscience, it is perhaps unsurprising that most studies of resting and task-evoked FC (rFC and tFC, respectively) focus on communities that are assortative in character.

Fundamentally, however, networks can exhibit diverse community structure. Of course, some communities can be assortative, but others can contain nodes that interact in non-assortative configurations. The two most commonly studied examples of non-assortative communities are (i) core-periphery structures, in which a densely connected core projects to a sparsely connected periphery, and (ii) *dis*assortative structures, in which the strongest connections fall between communities. Whereas assortative community structure emphasizes the segregative features of a network, non-assortative communities emphasize a network’s integrative properties and reflect its capacity for communication and signalling across community boundaries (Fig. 1). Core-periphery structure is thought to support a core’s ability to transiently broadcast information to or receive information from the periphery, while disassorative structure is an effective architecture for information transmission between two populations of computational units. Non-assortative communities are not easily dismissed as purely theoretical constructs; in fact, they are common in many biological and socio-technological networks [22–28], in which they occupy unique functional roles. Despite this evidence, there are few examples in the network neuroscience literature in which non-assortative community structure has been explored in earnest [25, 29]. Most studies continue to use better-established (though perhaps limiting) methods for community detection. In doing so, these studies remain insensitive to the potential richness and diversity of community structure not captured by modularity maximization or Infomap, and in some cases run the risk of fundamentally mischaracterizing that structure.

Collectively, these observations, together with recent theoretical work showing that real-world networks may have no uniquely optimal community structure [30], motivate the application of new and unexplored methods for uncovering communities in rFC and tFC networks. Here, we address these challenges directly using a weighted stochastic blockmodel (WSBM) [31, 32] to detect both assortative and non-assortative community structure in FC data taken from the Human Connectome Project (HCP) [33]. In line with current thinking, we find that rFC is dominated by assortative communities. However, we also find evidence for non-assortative organization in higher-order association areas, suggesting that the polyfunctionality of these areas may be underpinned by their deviations from pure assortativity. Next, we show that compared to rFC, tFC is accompanied by a reduction in assortativity, offset by increases in the prevelance of core, periphery, and disassortative communities. These findings suggest that cognitively demanding tasks are not simply a tuning of connection weights among a fixed set of assortative communities, but a wholesale reconfiguration of the brain’s functional architecture into novel non-assortative structures supportive of information transmission. Next, we show that network measures derived from communities facilitate the accurate classification of which task a subject is performing. Importantly, network measures of non-assortativity outperformed those associated with assortativity. Finally, we show that intersubject differences in behavioral measures are correlated with non-assortative features of brain areas, and that these correlation patterns are unique across tasks but nonetheless show high specificity to intrinsic functional systems. These findings offer a new perspective on the brain’s community organization and its relation to cognition.

## RESULTS

In this report we use the WSBM to detect communities in rFC and tFC (see **Materials and Methods** for details of WSBM implementation and network construction). Here, we describe the results of those analyses, broken down into four sub-sections. First, we characterize the non-assortative community structure of rFC in the subsection entitled **Community structure at rest**. In the sub-section entitled **Task-based community structure is diverse and non-assortative**, we repeat the analyses carried out in the first section but for tFC estimated during seven different cognitive tasks and a second resting-state run. Next, we derive a set of area-level network measures based on community structure that we use as features for training a random forest classifier. In the subsection entitled **Classifying task/cognitive state**, we show that these measures, in particular those based on non-assortative communities, can accurately classify the task state of held-out subjects. Finally, in the subsection entitled **Non-assortative community structure is correlated with cognitive performance**, we show that inter-subject differences in these same network measures are correlated with subject performance on cognitively demanding tasks.

### Community structure at rest

Past studies have described rFC community structure as uniformly assortative: strong FC exists on edges that are distributed densely within communities but sparsely between communities. This description belies the diversity and range of configurations that networks and their communities can adopt, including core-periphery and dis-assortative configurations (see Fig. 2 for illustrated examples). Here, we define four network measures that index each brain area’s participation in assortative, core, periphery, and disassortative communities. We define two additional measures that quantify the diversity of communities that areas participate in as well as their dominant community class (see **Methods and Materials** for more details). We calculate these measures for individual subjects and present their average values. We report results for simulations in which the number of communities is fixed at *k* = 6, and we show that these results hold for other values of *k* in Fig. 3m.

In general across the rFC data, we find that brain areas predominantly participate in assortative communities, in agreement with extant literature (Fig. 3a,g). The most assortative brain areas include those traditionally associated with primary sensory systems supporting visual and somatomotor function, as well as those associated with attentional and salience systems supporting higher-order cognitive functions. A number of brain areas show markedly lower assortativity relative to others; these include areas traditionally assigned to the default mode and cognitive control systems (Fig. 3). The explanation for these areas’ reduced assortativity is evident when we examine the areas that are most core-like or most periphery-like, which are near-perfect complements of the assortative map (Fig. 3b,h and Fig. 3c,i). We find that these less assortative areas exhibit inter-digitation, with peripheral areas spatially surrounding core areas. These findings dually support and challenge our current understanding of the brain’s community structure. In agreement with past studies, we find that most communities are assortative. However, we also find non-trivial levels of non-assortative communities in the form of core-periphery motifs. Interestingly, the areas with the greatest expression of cores and peripheries map onto the brain’s default mode and cognitive control networks, both of which are consistently associated with higher-order cognitive processing [34], suggesting a neuro-topological explanation for these areas’ functionality.

**FIG. 1.**
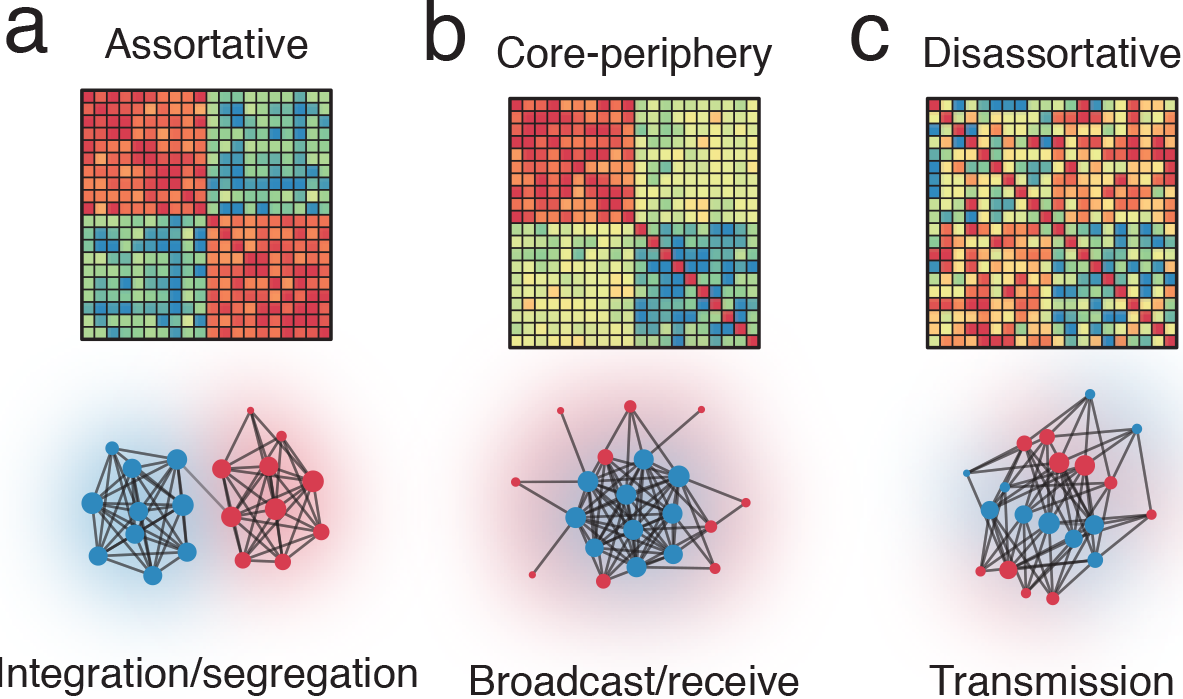
Schematic illustrating assortative and non-assortative community structure. Different classes of communities reflect and emphasize different communication policies. (*a*) Assortative communities, for example, highlight the segregation of information between communities and the diminished capacity for information to be integrated across the boundaries of communities. (*b*) In contrast, core-periphery structure highlights network structures that reflect the capacity for information to be broadcast from a core to a set of receivers, the periphery. (*c*) Finally, disassortativity reflects network structures that may be poised for transmitting information and signals away from other members of the same community.

Next we turned to a consideration of the diversity of communities in which each brain area participates. Intuitively, we would consider a brain area that disproportionately expresses a single class of community to be less diverse than a brain area that expresses many different classes. We find that this diversity measure varies across the cortex, peaking in the same areas that participate in core-like and periphery-like communities: areas traditionally associated with default mode, cognitive control, and salience systems (Fig. 3e,k). To better understand an area’s preference for community structure, we assigned each brain area a dominant community class by *z*-scoring each class’ brain-wide expression values and, for each brain area, identifying the class associated with the greatest *z*-score. As expected, community dominance maps revealed that primary sensory systems are dominated by assortative communities while core and periphery classes were concentrated within default mode and cognitive control systems (Fig. 3f,l). Importantly, we show in the supplement that these brain-wide patterns are reproducible; the expression of assortative, core, periphery, and disassortative communities as well as the diversity measures are highly correlated across two separately-acquired resting-state scans (Supplementary Fig. S1).

Collectively, these findings offer new insight into the brain’s meso-scale organization at rest, challenging current accounts in which communities are assumed to be uniformly assortative. Our findings paint a picture in which functional network community structure is fundamentally diverse – a feature that may support complex cognitive processes [29, 35]. Interestingly, the differential expression of assortative *versus* non-assortative communities is greatest when comparing primary sensory systems (almost entirely assortative) and the default mode system (expresses cores and peripheries). This observation aligns with the intuition that segregated processing is a key feature supporting specialized brain function, while poly-functionality is a property that requires intercommunity interactions and communication.

**FIG. 2.**
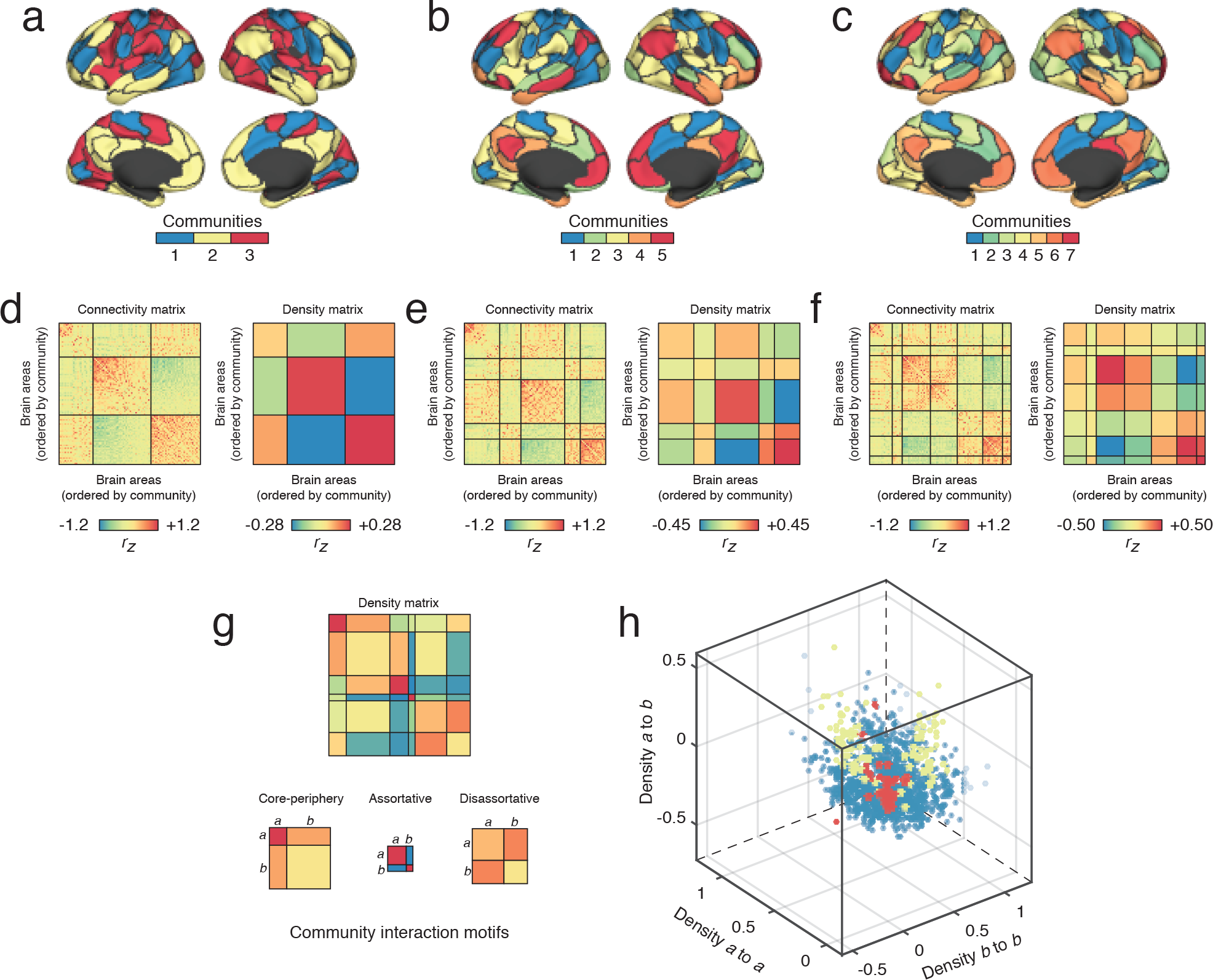
Typical WSBM output and schematic. We show examples of communities detected with the number of communities set to *k* = 3, 5, 7. Panels (*a*)-(*c*) depict topographic representations of community assignments for given brain areas. Panels (*d*)-(*f*) depict adjacency and connection density matrices with rows and columns ordered by areas’ community assignments. (*g*,*h*) Based on intra- and inter-community connection densities, we can situate every pair of communities in a three-dimensional morphospace. This space can be partitioned so that each point, representing pairs of communities, is uniquely labeled as either an assortative, disassortative, or core-periphery interaction.

### Task-based community structure is diverse and non-assortative

Next, we fit the WSBM to tFC generated from data acquired while subjects performed cognitively demanding tasks inside the MRI scanner. We refer to these tasks as EMOTION, GAMBLING, LANGUAGE, MOTOR, RELATIONAL, SOCIAL, and WM (working memory) (see **Materials and Methods** for details on the specific tasks). As in the previous section, we calculated area-level measures that quantify the expression of assortative, disassortative, core, and periphery communities in addition to the diversity of community expression. We compared these measures across tasks and against both resting state scans.

Broadly, we observed that community expression patterns were only weakly correlated across tasks (Fig. 4a). That is, communities estimated from tFC networks were less segregated from one another, on average, compared to resting-state conditions (non-parametric test in which task labels were permuted randomly and independently for each subject; *p* < 0.01). We also found that tFC was associated with decreased assortativity (Fig. 4b) and increased community diversity (*p* < 0.01; Fig. 4c), as well as a proportional increase in core, periphery, and disas-sortative interactions (Fig. 4d-f). Importantly, the overall reduction in assortativity and the extent to which it was replaced by non-assortative interactions varied as-cross tasks, with RELATIONAL and GAMBLING tasks corresponding to the greatest and slightest reductions in mean assortativity, respectively. In Supplementary Fig. S2, we show that these general results hold as we vary the number of detected communities, *k*.

**FIG. 3.**
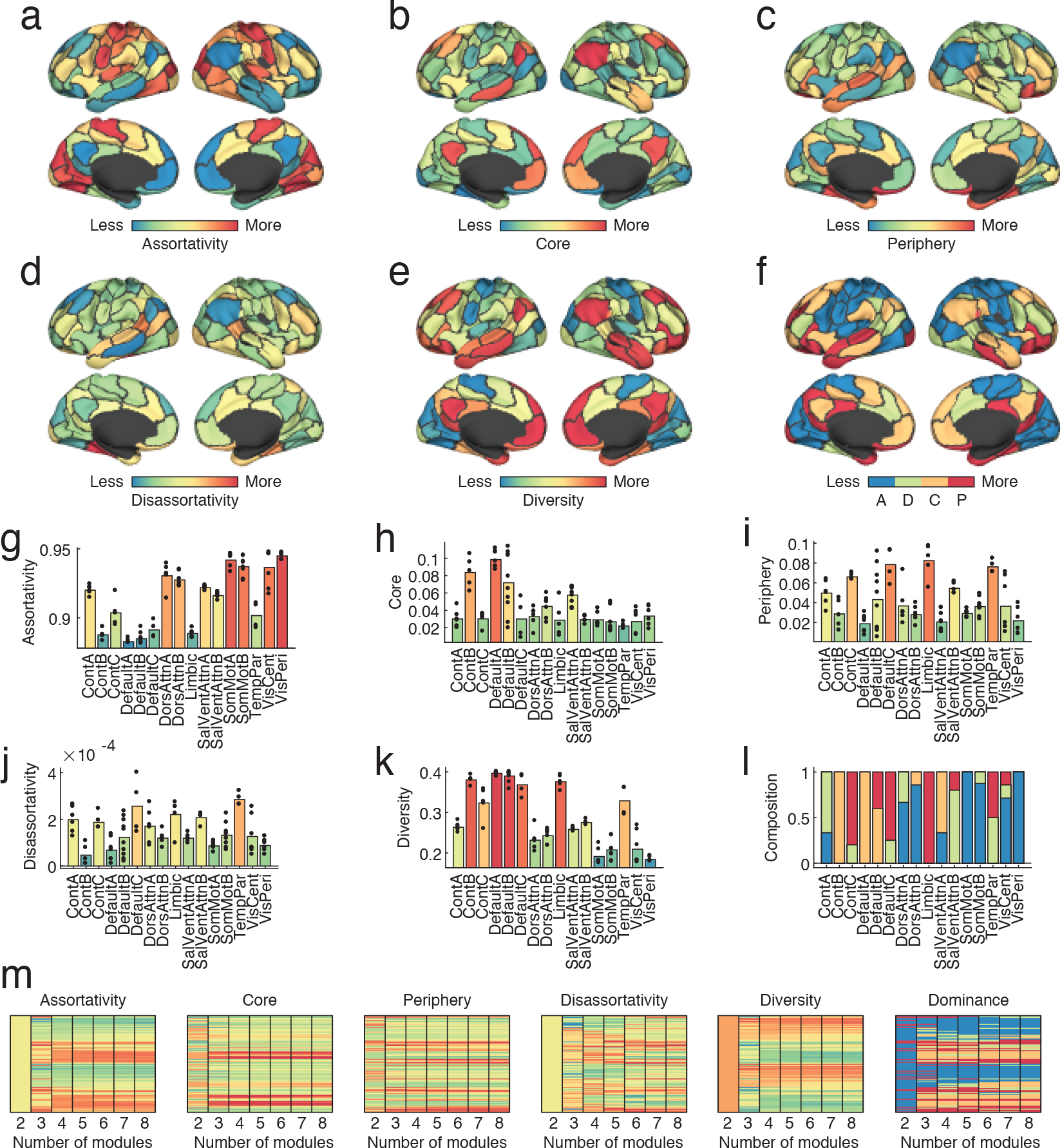
Group-level non-assortative community structure at rest. We averaged community participation, diversity, and community class dominance scores across subjects in order to characterize the brain’s community structure at the group level. Here we show topographic representations of brain areas’ participation in assortative (*a*), core (*b*), periphery (*c*), and disassortative (*d*) interactions. Panels (*e*) and (*f*) depict brain areas’ diversity and community dominance scores (‘A’ for assortativity, ‘D’ for disassortativity, ‘C’ for core, and ‘P’ for periphery). Panels (*g*) - (*l*) depict the same data as shown in the brain plots but now aggregated by brain systems. The number of communities is fixed at *k* = 6 for panels (*a*) through (*l*). In panel (*m*) we show the consistency of the above results as a function of the number of detected communities. Assortativity, core, periphery, community diversity, and community dominance scores change very little. The disassortativity score is more variable and present at low levels. Note: The color of bars in panels (*g*) through (*m*) denote system-averaged values.

These findings broadly agree with past reports documenting tFC tradeoffs in community segregation and integration [10, 14] and strengthened between-community connections in tFC compared to rFC [36]. While these past studies arrive at such conclusions after assuming that the brain is composed of strictly assortative communities, our study suggests that increased integration is a consequence of fundamental changes in the character of communities and the increased prevalence of cores and peripheries. That is, task-based reconfiguration of FC is driven less by subtle fluctuations in the connection weights between assortative communities, and instead by the emergence of novel non-assortative communities. These findings suggest that non-assortativity in community structure may be a reflection of ongoing cognitive processes.

**FIG. 4.**
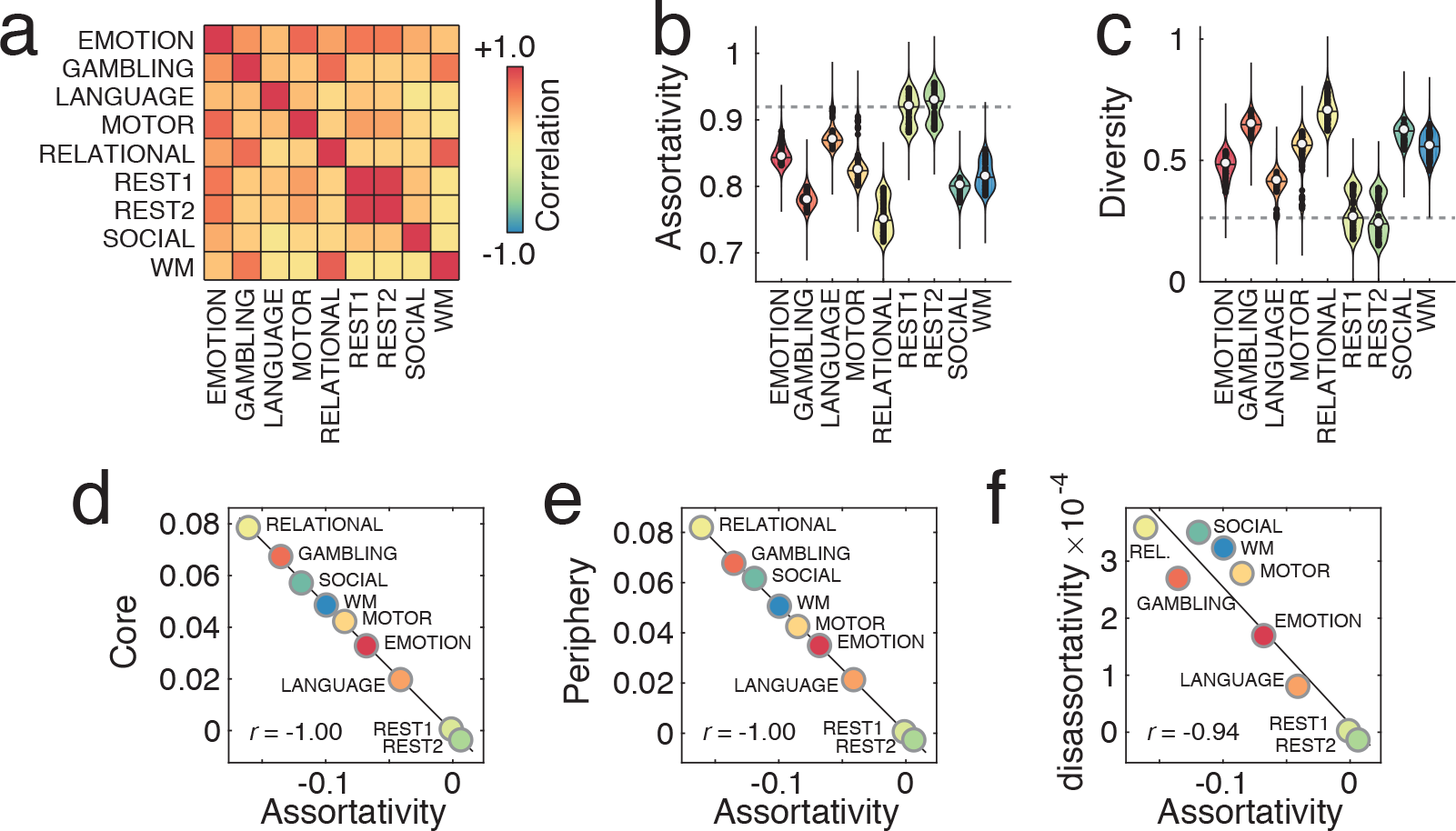
Task-evoked FC and community structure. For each task and for each brain area, we averaged expression levels of assortativity, disassortativity, core, and periphery communities across subjects. (*a*) We assembled these mean expression levels into a single vector for each task, and for every pair of tasks we computed the correlation between their corresponding vectors. (*b*) Violin plots of areal assortativity expression levels. Note that both resting state runs achieve the greatest mean assortativity (REST1 and REST2), while assortativity levels for tasks are consistently and statistically lower. (*c*) Violin plots of the community diversity measure. We see that resting state runs achieve the lowest diversity levels. (*d,e,f*) Scatterplots depicting association of average core, periphery, and disassortativity levels as a function of average assortativity.

### Classifying task state

In the previous section we showed that tFC is characterized by decreased assortativity as the prevelance of core, periphery, and disassortative communities increases. The fact that these changes were, on average, uncorrelated across tasks suggests the possibility of classifying a subject’s cognitive state as operationalized by the task that they are performing inside of the scanner. In this section, we test this possibility directly and train random forests composed of 100 weak classifiers to classify tasks using as features brain areas’ assortative, dis-assortative, core, and periphery expression levels. To reduce the possibility of overfitting, we adopted a cross validation strategy in which random forest classifiers trained on data from 90% of the subjects were used to predict the task states of the held out 10% (25 repetitions) (See **Materials and Methods** for more details).

We explored two strategies for choosing features. The first strategy allowed us to determine which brain areas were most useful for the classification of task state. Specifically, we defined for each brain area a five-dimensional feature vector composed of its assortative, disassortative, core, and periphery expression levels in addition to its diversity index. We then looped over brain areas, and at each iteration we used features from a single brain area to train the classifier. In the second strategy, we performed principal component analyses on the *N* × *N*_*subject*_ matrices of community expression levels and the diversity index. For each measure, we treated the first *P* principal component coefficients as features that were then used to train the classifier.

#### Predicting task state with single-area features

We first tested whether features of individual brain areas could be used to correctly classify the subjects’ task states (see Fig. 5a for a schematic). In general, we found that the classifier’s mean true positive rate (TPR) was above chance for all tasks, but was especially adept at differentiating the resting state from all task states (Fig. 5b). To visualize these results, we plotted task-averaged TPR on the cortical surface (Fig. 5c). We found that TPR was greatest in brain areas associated with the visual and dorsal attention systems (Fig. 5d). In some ways, this observation is unsurprising. Given that the classifier most accurately classified the resting state scan and that rFC was characterized by high levels of assortativity in dorsal attention, motor, and visual areas, we might expect that the classifier’s overall performance would be driven by these same areas.

**FIG. 5.**
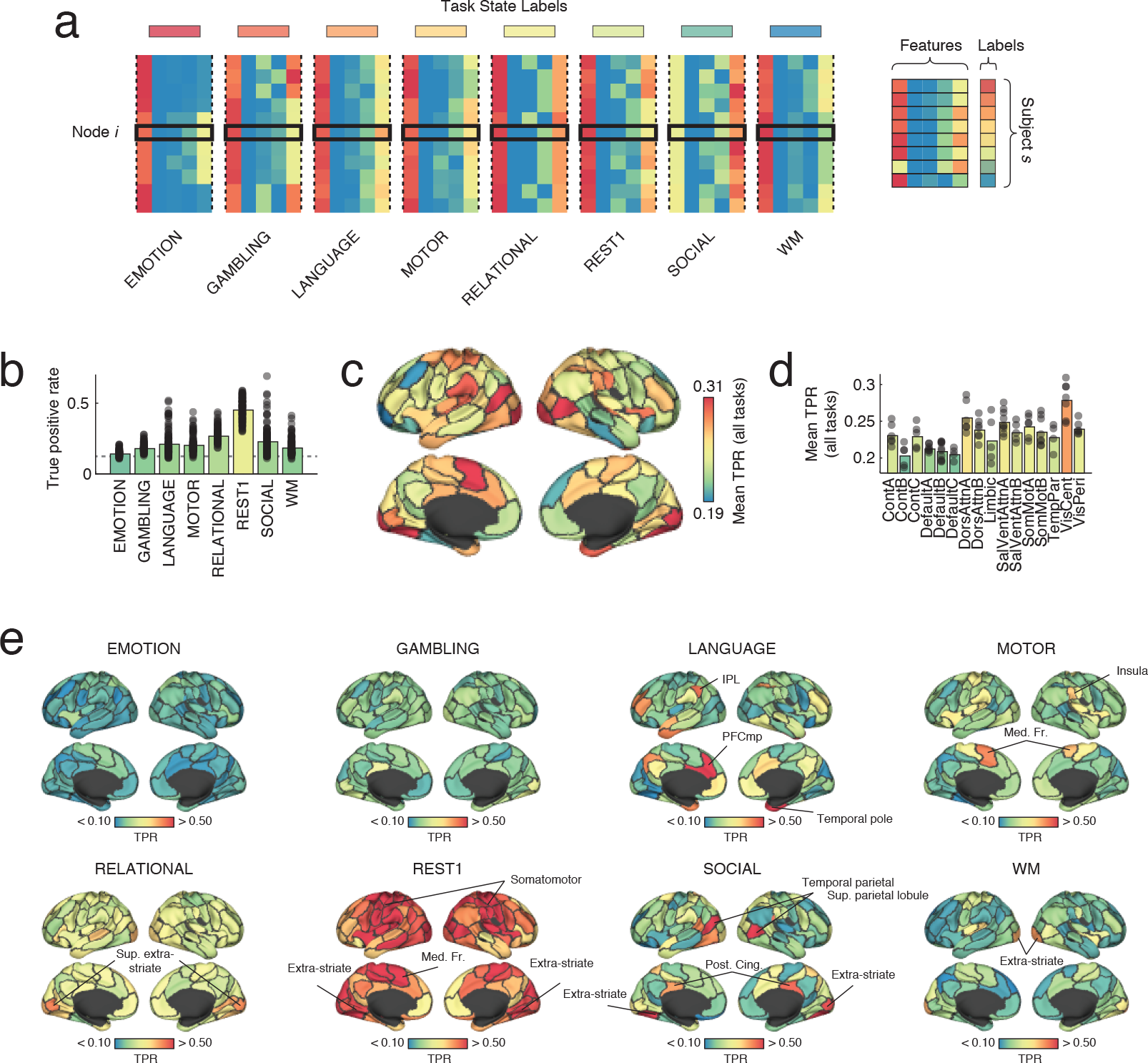
Classification using areal features. We used random forest classifiers to predict task state as operationalized by the scan from which the data was taken. (*a*) For each brain area and for each task state (here depicted as colored bars), we defined a five-dimensional set of features based on that area’s assortative, disassortative, core, and periphery expression levels in addition to its community diversity score. These features were used to train a random forest classifier and predict the task state of held-out data. (*b*) We found that mean performance across areas was greater than chance for all task states. Overall, classification was greatest for the resting-state run. (*c,d*) The areas with the greatest true positive rate across all tasks were located in visual and dorsal attention systems. (*e*) Areas were differentially effective in classifying particular task states. Note: The color of bars in panels (*b*)and (*e*) denote the mean TPR over all runs of the classifier.

We also examined brain-wide TPR for each task independently and found considerable heterogeneity in the brain areas that were useful for correctly classifying different tasks. For instance, while most brain areas exhibited low TPR when classifying the LANGUAGE task, medial prefrontal cortex, temporal pole, and inferior parietal lobule all exhibited true positive rates greater than 50%. Similarly for the SOCIAL task, we found that superior parietal lobule along with extra-striate and temporal parietal cortices exhibited especially high TPR. In the supplement, we show that we found similar spatial patterns when we varied the number of detected communities (Supplementary Fig. S3).

These findings suggest that variation in the expression levels of community classes at individual brain areas carries information about an individual’s task state and by extension the cognitive processes elicited. Interestingly, and with the exception of rFC, we also found that for a given task only a small subset of brain areas exhibited above-chance true positive rates, suggesting that the capacity to discriminate between task states is not a brainwide phenomenon, but is instead limited to specific sub-networks and anatomical locations.

#### Predicting task state with whole-brain features

In the previous section we used features defined at the level of individual brain areas to train and classify subjects’ task states. In this section, we use low-dimensional projections of whole-brain features in the form of principal component coefficients to solve the same classification problem. Specifically, we tested five raw feature sets: expression levels of assortative, disassortative, core, and periphery communities as well as community diversity scores (Fig. 6a). For each feature set, we pooled its values across all tasks and performed a principal component analysis (Fig. 6b). This analysis resulted in a set of orthonormal principal component (PC) scores that served as a basis set. Each score was accompanied by a set of PC coefficients, which represented the projection of subject-level data onto each PC score. Upon visual inspection we found that tasks were sometimes segregated from one another in PC space, where locations were defined based on PC coefficients (Fig. 6c).

**FIG. 6.**
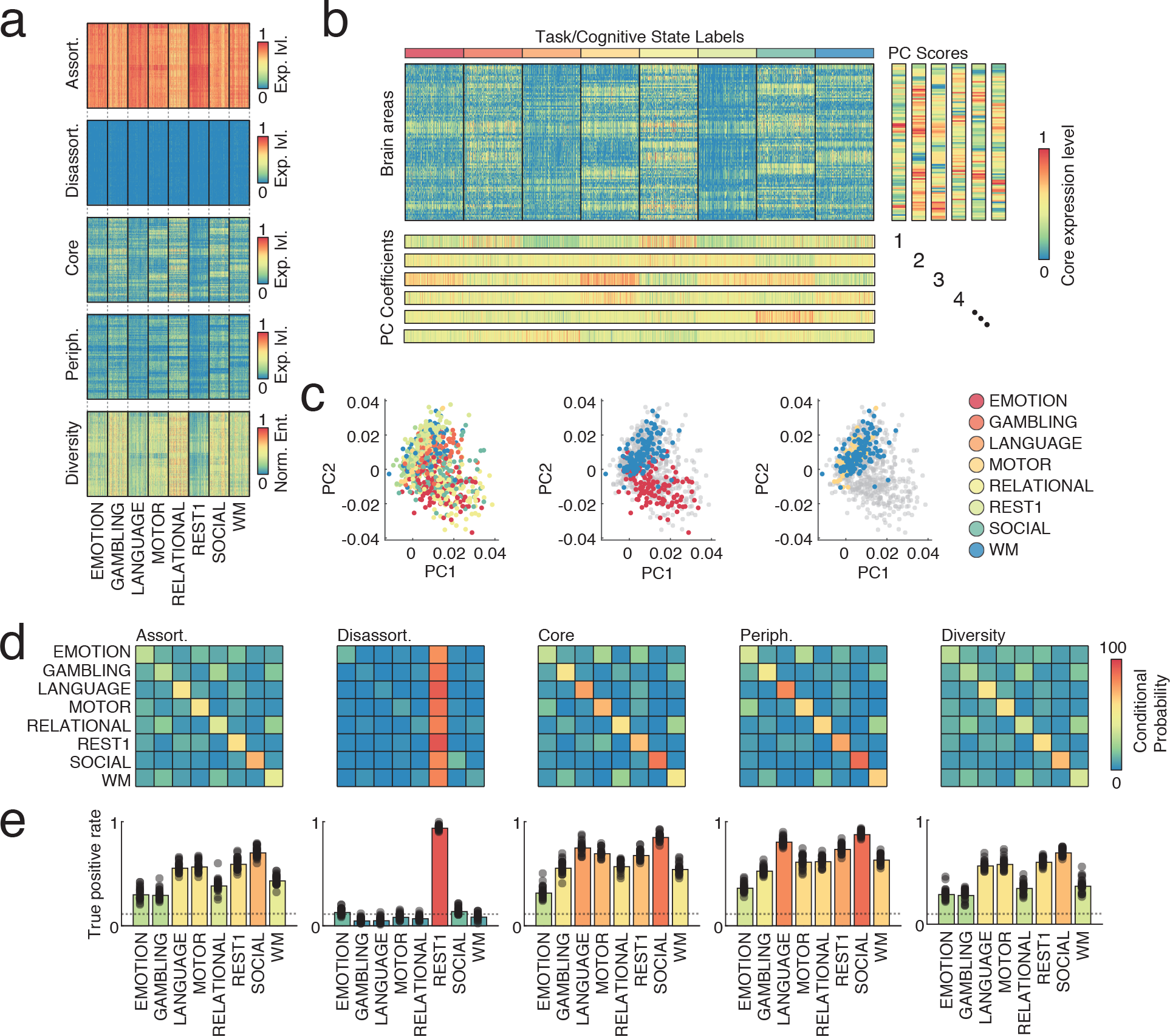
Classification using brain-wide features. We used random forest classifiers to predict scan type. (*a*) We tested five sets of raw features. Each subject and each scan was characterized by a brain-wide pattern of assortativity, disassortativity, core, and periphery expression levels in addition to the community diversity score. (*b*) For each of these feature sets, we performed a principal components analysis, which generated an orthonormal set of principal component scores along with coefficients representing projections of the brain-wide features into a lower-dimensional feature space (see, for example, panel (*c*)). Coefficients were repeatedly divided into 90/10 training/test splits. We used the first *P* components as inputs to the random forest model, and then used the fitted model to predict the task state of the held-out data. Panels (*d*) and (*e*) depict results with the number of communities and features both fixed at *k* = *P* = 6. (*d*) Confusion matrices generated from classifier output. Within each row, the value of the cell indicates the probability that specific task states were correctly classified. (*e*) For each feature type, we show the true positive rate for correctly classifying each of the seven task states and the resting state. Note: The color of bars in panel (*e*) denote the mean TPR over all runs of the classifier.

Next, we trained random forest classifiers using the first *P* PC coefficients as features. Suprisingly, we found that even a small number of PC coefficients were sufficient for generating high classification accuracy. Interestingly, the exact level of accuracy varied by feature type and by task. With the exception of disassortativty, which showed particularly poor specificity, all features outperformed chance by a wide margin (Fig. 6d,e). Across features, SOCIAL, LANGUAGE, and REST were among the tasks that were most frequently classified correctly. Interestingly, the features corresponding to the greatest overall TPR were core and periphery expression levels (0.62 and 0.65, respectively; compared to 0.48, 0.19, and 0.46 for assortative expression level, disassortative expression level, and community diversity score, respectively). In Supplementary Fig. S4, we show that these reported results are consistent as we vary the number of detected communities, *k*, and the number of principal components, *P*, used by the random forest model to perform the classification.

These findings demonstrate concretely that measures of non-assortative community structure, namely the expression levels of cores and peripheries, are especially useful for accurate classification of subjects’ task states. Importantly, Infomap and modularity maximization seek only assortative communities, and thus the communities that they identify might not be optimally suited for task state classification. Our findings motivate the future consideration of non-assortative communities and the WSBM in a translational context, for example to distinguish neural phenotypes in patients versus controls [37–39].

### Non-assortative community structure is correlated with cognitive performance

In the previous sections we characterized community structure in both rFC and tFC, and we demonstrated that under certain conditions measures of non-assortativity could be used to correctly classify subjects’ task states. An important remaining question is whether measures of non-assortative community structure are associated with subject performance on cognitively demanding tasks. Here we address this question by computing correlations of brain areas’ community expression levels and subject performance on MEMORY, SOCIAL, RELATIONAL, and LANGUAGE tasks (see **Materials and Methods** for information on additional pre-processing of the behavioral data). To simplify our analyses and shift focus onto brain systems, we subsequently computed system-level correlations by averaging over cognitive systems’ constituent areas. We expressed these system-level correlations as *z*-scores relative to a null distribution generated by randomly and uniformly permuting areas’ system labels (1000 repetitions). Additionally, we limited the scope of our analyses focusing exclusively on rFC, as there exists an extensive body of prior work focused on behavioral correlates of other measures computed on rFC data.

We calculated Spearman correlations for all pairs of brain and behavioral measures and averaged these coefficients at the level of brain systems. Interestingly, correlation patterns were largely unique to each brain-behavior combination, which we can visualize by embedding patterns in two-dimensional space using multi-dimensional scaling (Fig. 7a,b). Here, we find no obvious clustering: correlation patterns are not co-localized according to behavior or brain measure, suggesting that the expression levels of each community class offers a complementary perspective on the relationship between brain networks and behavior.

**FIG. 7.**
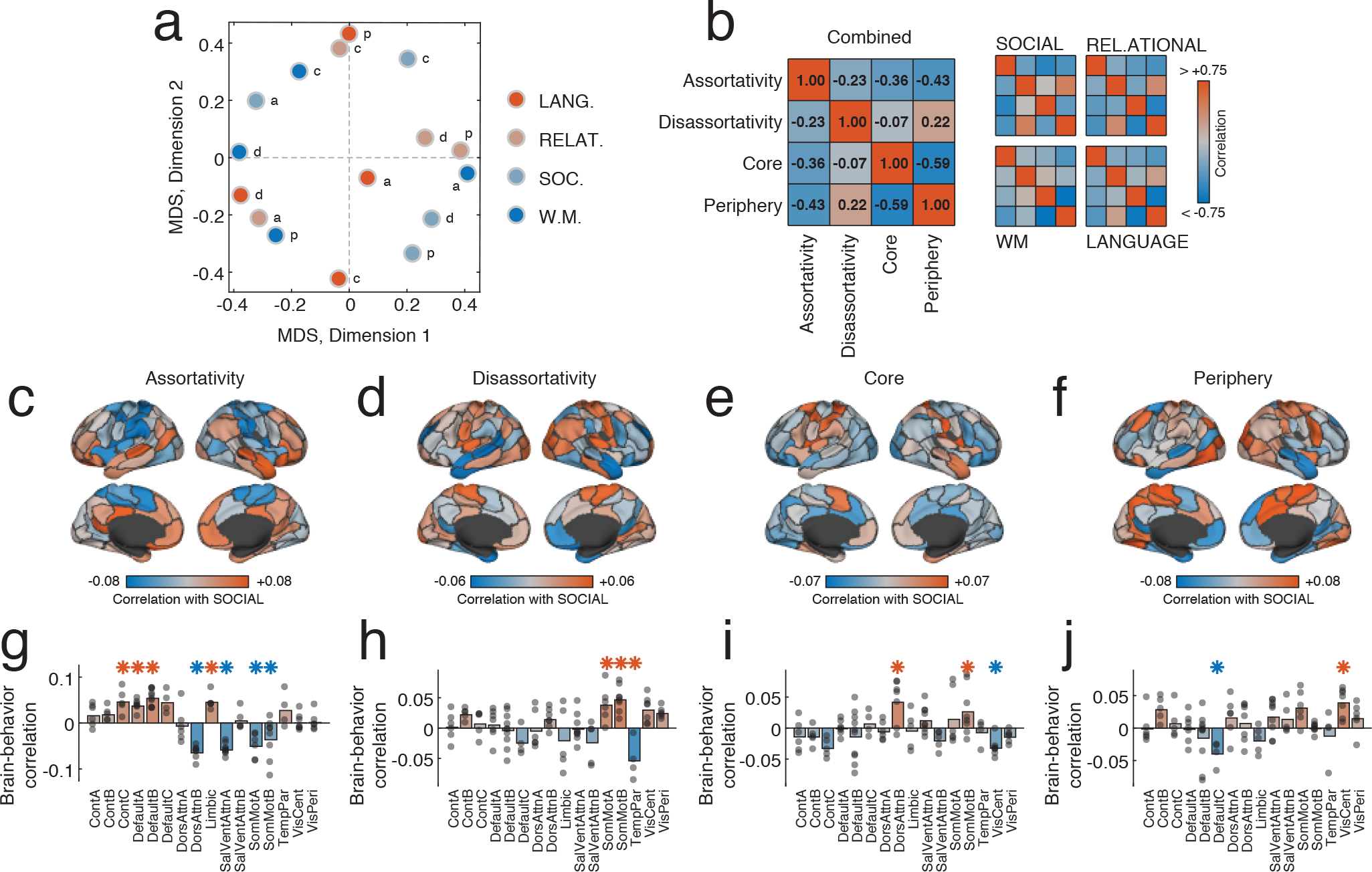
Correlations of community features with behavioral measures. (*a*) For each network measure - assortativity, core, periphery, and disassortativity – we generated a *N* × 1 vector of correlation coefficients. We show, here, the correlation pattern among those vectors for all behavioral measures combined and for each behavioral measure independently. In general, we find that behavioral correlations are dissimilar across measures, with the strongest dissimilarity between core and periphery correlations. (*b*) We can visualize this result by using multidimensional scaling to embed each correlation pattern in twodimensional space while approximately preserving inter-pattern distances (correlations). In general, we find that patterns from the same behavioral measure are not located near one another. Correlation of subjects’ performances on a SOCIAL cognitive task with their (*c*) assortativity, (*d*) disassortativity, (*e*) core, and (*f*) periphery scores. In panels (*g-j*) we aggregate those scores by brain systems. Red and blue asterisks indicate system-level correlations that were more positive or negative than chance; *p* < 0.05 while fixing the false discovery rate (FDR) at 5%. In panels *c-j*, colorbar indicates magnitude of correlation.

Next, we investigated the area- and system-level correlation patterns in greater detail. Here, we show correlations of the SOCIAL behavioral measure with assortativity, disassortativity, core, and periphery expression levels. Though the overall correlation magnitudes are relatively weak, we find surprisingly strong system-level effects. For instance, the correlation of assortativity expression levels with the SOCIAL measure is consistently positive within the control and default mode systems but negative in the attention and motor systems (*p* < 0.05, FDR-controlled; Fig. 7c,g). We find similar results for disassortativity (*p* < 0.05, FDR-controlled; Fig. 7d,h; positive correlations in somatomotor systems; anticorrelations in temporal parietal system), core (*p* < 0.05, FDR-controlled; Fig. 7e,i; positive correlations in thedorsal attention system), and periphery (*p* < 0.05, FDR-controlled; Fig. 7f,j; positive correlations in the visual and somatomotor systems; negative correlations in the default mode system). We show analogous correlation maps for WORKING MEMORY, RELATIONAL, and LANGUAGE measures in Supplementary Fig. S5 and their variation with *k*, the number of communities, in Supplementary Fig. S6.

These findings suggest that cognitive performance, even when assayed against the brain at rest, is modestly correlated with the expression levels across different classes of community structure. The overall magnitude of these correlations is weak, but their areal and system-level distributions are highly specific, concentrating within particular cognitive systems. Interestingly, in the context of the SOCIAL task, we find the assortativity of default mode components to be correlated with performance, which agrees with past studies suggesting that this system may play an important role in social cognition, mentalizing, and theory of mind [40]. Our findings also implicate the somatomotor network, whose assortativity, disassortativity, and coreness are all significantly correlated (or anti-correlated) with performance on the SOCIAL task. This observation is consistent with recent findings that link motor behavior and social cognition [41, 42]. More generally, however, the topography of correlation patterns is specific to each pair of behavioral and network measures, suggesting that different aspects of community structure encode unique information about brain-behavior associations. In future studies, these measures might prove useful in generating biomarkers for clinical conditions by testing for associations with continuously defined clinical measures [43], or in identifying developmental and aging phenotypes by testing for associations with age [44].

## DISCUSSION

In this report we leverage recent methodological advances in community detection algorithms to resolve both assortative and non-assortative communities in resting and task-evoked FC. We find that rFC is characterized by high levels of assortativity, with the most assortative brains areas corresponding to primary sensory systems and the least assortative brain areas corresponding to default mode and control systems. We next show that tFC is less assortative than rFC, displaying an increased prevelance of core and periphery communities. We show that these differences are task-specific and can be used to classify subjects’ task state with accuracies greater than chance. Finally, we show that inter-individual differences in community structure are correlated with subjects’ performances on in-scanner tasks. These findings present an alternative perspective on task-based reconfiguration of functional brain networks, and open up avenues for future studies to apply blockmodeling techniques to the study of development and aging, psychiatric disorders, and neurological disease.

### Resting-state functional network communities are not strictly assortative

Here we use the WSBM to uncover generalized community structure in rFC and find evidence that not all communities exhibit the stereotypical and expected internally dense, externally sparse organization. The presence of such non-assortative communities is unanticipated by past studies [5, 7, 45] likely due to the methodological biases of modularity maximization and Infomap, both of which are designed to detect assortative structure alone.

Our findings challenge the view that the resting brain is uniformly organized into segregated functional communities, which are thought to reflect specialized information processing. Rather, our findings suggest that the processing patterns of some brain areas may be nonspecialized and integrative, involving cross-talk between multiple systems and communities.

It is particularly notable that the brain areas with the greatest deviation from uniform assortativity are concentrated within default mode and cognitive control systems. Areas within these systems are generally considered to be poly-functional and supportive of many diverse cognitive processes [34, 46–48]. The non-assortativity of these areas together with the much greater assortativity in primary sensory areas suggests that non-assortativity may be an important determinant of an area’s functionality. Intuitively, greater non-assortativity may facilitate more integrative processing, thereby supporting higher order cognition. This hypothesis could be more carefully and systematically tested in future studies by, for example, using multi-task paradigms that tax different cognitive systems and comparing subjects’ performances to their non-assortative community expression patterns [49].

### Non-assortative community structure in tFC

While rFC exhibited modest yet areal-specific deviations from purely assortative community structure, we found that tFC exhibited a much greater shift towards non-assortativity. This observation is consistent with the hypothesis that cognitively demanding tasks recruit multiple brain systems and are underpinned by their transient communication with one another [11–13]. Also consistent with this hypothesis is the fact that these patterns of recruitment, which we characterize as reconfigurations of the brain’s community structure, are task-specific and aid in the accurate classification of individuals’ task states. Importantly, our findings show that classifications based on non-assortative structure (core and periphery expression levels) outperform those based on assortative structure, suggesting that non-assortativity plays a critical task-specific role.

Past studies have also described changes in FC patterns during cognitively demanding tasks, though these changes have always been expressed in terms of uniformly assortative community structure. That is, tFC may exhibit different community structure compared to rFC, but those communities are always assortative and cohesive [9, 10]. In contrast, we propose that tFC is characterized by the emergence of altogether novel classes of community structure and that where (in the brain) these communities form are largely specific to a given task. Collectively, our findings offer a novel perspective on task-based reconfiguration of community structure and, in characterizing that structure using newly-developed statistics [29], open avenues for future research [50]. Given the capacity of non-assortative communities for classification of task states, we hypothesize that similar measures could be applied to offer improved classification of neurodegeneration disease [51] and psychopathology [43], as well as to monitor the growth and development of neural circuits [44].

### Future work and extensions

Importantly, together with other recent papers [25, 29] the results reported here suggest that the traditional view of internally dense and externally sparse brain network modules is not privileged, and that newer and perhaps more general methods of community detection can offer complementary views. The implications of these findings are multifold. On one hand they serve a prospective role in which future studies can leverage a new suite of network summary statistics based on non-assortative communities that could be useful for biomarker generation and classification. Our findings also serve a role in retrospect, prompting us to reconsider results reported in past studies that were based on less general community detection tools like modularity maximization and Infomap.

Indeed, SBMs are becoming more common in network neuroscience. Recent applications have demonstrated that non-assortative community detection methods applied to structural connectivity can better reproduce patterns of FC and correlated gene expression compared to traditional methods [29]. Other studies have further demonstrated that annotated SBMs can be used to constrain community structure based on patterns of neural activity [52], and that SBMs naturally detect higher-order network structures like rich clubs [25]. These applications mirror a more general trend in the network science community where increased emphasis is being placed on developing SBM-related methods for weighted and hierarchical networks [53, 54], multi-layer networks [55, 56], and annotated graphs [57].

Additionally, recent results have shown that for real-world networks where the ground truth community structure is unknown, there exists no uniquely optimal community detection algorithm [30]. This observation further motivates the exploration of a plurality of different community detection tools on any given dataset, with each method providing potentially complementary perspectives. The continued reliance on Infomap and modularity maximization in the still-young network neuroscience community may actually have a limiting effect by distracting from or obscuring potentially important features of brain networks. A less parsimonious but more fruitful strategy moving forward would be to seek both the points of convergence and divergence across techniques and to tease out their unique contributions.

### Limitations

This study is limited in several ways. First, just as Infomap and modularity maximization have known biases and shortcomings, so too do blockmodels. Among those is the so-called “detectability limit” [58, 59] that prevents the detection of true communities in networks whose sparsity is below some critical value. Note that this phenomenon is analogous in many ways to the resolution limit associated with some versions of the modularity quality function [60]. Exactly how this limit scales to the WSBM or fully-weighted matrices is unclear, but it may play an important role in the WSBM’s ability to detect certain classes of communities.

Another potential limitation is the use of correlation as a measure of FC. In practice, individual functional connections are not independent of one another [61], an observation that has been shown to artificially inflate certain network statistics. The WSBM, however, assumes that connections are drawn independently and randomly from a given distribution. Though SBMs are routinely applied to other networks whose connection weights exhibit similar dependencies, future work could be directed to developing generative models with parameterized block community structure that build in assumptions appropriate for correlation (or equivalently covariance) matrices.

Another issue concerns the absence of sub-cortical structures in our analysis. Though the choice to exclude these areas is in line with other studies of community structure in FC [5], sub-cortical structures are known to play key regulatory and functional roles in neural dynamics and cognition. Again, future studies could be designed that explicitly probe the non-assortative structure of sub-cortical FC and its interactions with the cerebral cortex [62, 63].

A final limitation concerns how we interpret FC. At times we have treated functional connections as though they were pathways along which inter-areal communication takes place. In reality, of course, FC is simply the statistical similarity structure that emerges naturally as result of a dynamical system whose temporal evolution is constrained by anatomical connectivity along with genetic, spatial, and environmental factors [64–66]. As such, the communities that we detect reflect the outcome of an ongoing dynamical process rather than the process itself. Future work must jointly investigate the roles of structure and dynamics in shaping all aspects of FC, including its community structure. Neural mass models, for example, offer a neurobiologically plausible and mechanistic account of how FC patterns emerge and fluctuate [64, 67], while the growing field of network control describes the evolution of network neural systems in the presence of exogenous inputs [68–71].

## CONCLUSION

In summary, we show that both rFC and tFC exhibit elements of non-assortative community structure, challenging the widely held belief that brain network communities are uniformly assortative. We also demonstrate that measures derived from non-assortative communities, specifically, are useful for classifying subjects’ task states and are related to behavioral performance on cognitive tasks. The results reported here demonstrate the utility of blockmodels as an approach for studying meso-scale structure in FC and open up important avenues for future work.

## MATERIALS AND METHODS

### Human Connectome Dataset

In this study we aimed to characterize the assortative and non-assortative community structure of resting and task-evoked FC. To address this aim, we leveraged data from the Human Connectome Project (HCP), a multisite consortia that collected extensive MRI, behavioral, and demographic data from a large cohort of subjects (>1000) [33]. As part of the HCP protocol, subjects underwent two separate resting state scans along with seven task fMRI scans. All functional connectivity data analyzed in this report came from these scans and was part of the HCP S1200 release [33]. Subjects that completed both resting-state scans and all task scans were analyzed. We utilized a cortical parcellation that maximizes the similarity of functional connectivity within each parcel (*N* = 100 parcels) [72].

We preprocessed resting-state and task data using similar pipelines. For resting-state, the ICA-FIX resting-state data provided by the Human Connectome Project were utilized, which used ICA to remove nuisance and motion signals [73]. For task data, CompCor, with five components from the ventricles and white matter masks, was used to regress out nuisance signals from the time series. In addition, for the task data, the 12 detrended motion estimates provided by the Human Connectome Project were regressed out from the time series, the mean global signal was removed, and the time series was bandpass filtered from 0.009 to 0.08 Hz.

To reduce artifacts related to in-scanner head motion, frames with greater than 0.2 millimeters frame-wise displacement or a derivative root mean square above 75 were removed [74]. Subjects whose scans resulted in fewer than 50% of the total frames left were not analyzed further; a total of 827 subjects met this criteria for all resting-state and task scans.

For all scans, the MSMAII registration was used, and the mean time series of vertices on the cortical surface (fsL32K) in each parcel was calculated. The functional connectivity matrix for each subject was then calculated as the pairwise Pearson correlation (subsequently Fisher *z*-transformed) between times series of all nodes. Both left-right and right-left phase encoding directions scans were used, and the mean matrix across the two (task) or four (resting-state) scans was calculated.

#### Behavioral data

All cognitive and behavioral performance measures were chosen *a priori* based on their use in a previous study [16], in which they were shown to be highly correlated with functional network architecture [16]. Accordingly, all analyses in the present study were carried out using these four behavioral variables. In the working memory tasks (WM score), we used the mean accuracy across all n-back conditions (face, body, place, tool). In the relational task (RELATIONAL score), we used mean accuracy across both the matching and the relational conditions. For the language task (LANGUAGE score), we took the maximum difficulty level that the subject achieved across both the math and language conditions. We did not use accuracy, because the task varies in difficulty based on how well the subject is doing, making accuracy an inaccurate measure of performance for these tasks. For the social task (SOCIAL score), given that almost all subjects correctly identified the social interactions as social interactions, we used the percentage of correctly identified random interactions.

Rather than analyze these four variables directly, we wanted to analyze each variable’s unique contribution. Indeed, all four variables (WM, RELATIONAL, LANGUAGE, and SOCIAL scores) were correlated with one another. So that we could shift focus onto unique contributions of each variable, we analyzed each score’s residuals after partialing out the effect of all other scores. For example, we partialed out the effect of subjects’ RELATIONAL, LANGUAGE, and SOCIAL scores from their WM score. As a result, the behavioral variables used for all analyses reported in this study were uncorrelated with one another.

### Stochastic blockmodel

The stochastic blockmodel (SBM) seeks to partition a network’s nodes into *k* communities. Let *z*_*i*_ ∈ {1,…, *k*} indicate the community label of node *i*. Under the standard blockmodel, the probability that any two nodes, *i* and *j*, are connected to one another depends only on their community labels: 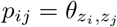.

To fit the blockmodel to observed data, one needs to estimate the parameters *θ*_*rs*_ for all pairs of communities {*r*, *s*} ∈ {1,…, *k*} and the community labels *z*_*i*_. Assuming that the placement of edges are independent of one another, the likelihood of a blockmodel having generated a network, *A*, can be written as:

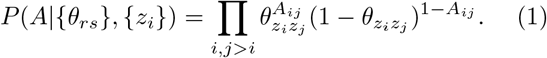

Fitting the SBM to an observed network involves selecting the parameters {*θ*_*rs*_} and {*z*_*i*_} so as to maximize this function.

### Weighted stochastic blockmodel

The classical SBM is most often applied to binary networks where edges carry no weights. In order to maximize its utility to the network neuroscience community where most networks are weighted, the SBM needs to be able to efficiently deal with weighted edges. Recently, the binary stochastic blockmodel was extended to weighted networks as the weighted stochastic blockmodel (WSBM) [32].

To understand the chosen form of the WSMB, we first note that Eq. 1 can be rewritten in the form of an exponential family of distributions [32]:

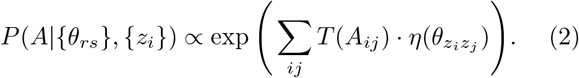

For the classical (unweighted) SBM, *T* is the sufficient statistic of the Bernoulli distribution and *η* is its function of natural parameters. Different choices of *T* and *η*, however, can allow edges and their weights to be drawn from other distributions. The WSBM, like the classical SBM, is parameterized by the set of community assignments, {*z*_*i*_}, and the parameters 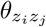. The only difference is that 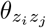 now specifies the parameters governing the weight distribtion of the edge, *z*_*j*_*z*_*j*_.

Here, we follow [32], and model edge weights under a normal distribution, whose sufficient statistics are *T* = (*x*, *x*^2^, 1) and natural parameters *η* = (*η*/*σ*^2^, −1/(2*σ*^2^), −*μ*^2^/(2*σ*^2^)). Under this distribution, the edge *z*_*i*_*z*_*j*_ is parameterized by its mean and variance, 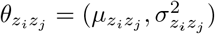, and the likelihood is given by:

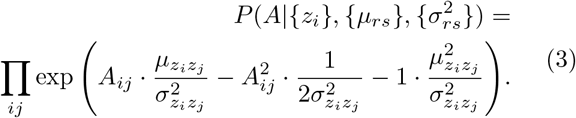

The above form assumes that all possible edges falling between communities are drawn from a normal distribution. The networks that we study here are fully-weighted, where a connection weight exists for all pairs of brain areas. In general, however, weighted networks may also be sparse, i.e. they may include edges where *A*_*ij*_ = 0 indicating the absence of a connection. The WSBM is designed with this general case in mind, but can easily accommodate fully-weighted networks. In the case of sparse networks, one can model edge weights with an exponential family distribution and model the presence or absence of edges by a Bernoulli distribution (akin to the unweighted SBM) [32]. Letting *T*_*e*_ and *η*_*e*_ represent the edge-existence distribution and *T*_*w*_ and *η*_*w*_ represent the normal distribution governing edge weights, we can rewrite the likelihood function for the sparse WSBM as:

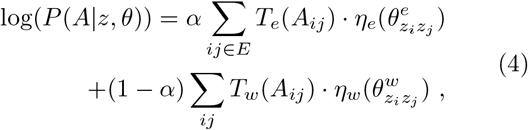

where *E* is the set of all possible edges, *W* is the set of weighted edges (*W* ⊂ *E*), and *α* ∈ [0,1] is a tuning parameter governing the relative importance of either edge weight or edge presence (or absence) for inference. In our application to fully-weighted FC networks, we fix *α* = 0. This effectively discounts any contribution from the binary graph’s topology so that the value of the likelihood function is determined only by contributions from the network’s edge weights.

For a given functional brain network, we maximize the likelihood of this sparse WSBM using a Variational Bayes technique described in [32] and implemented in MATLAB using code made available at the author’s personal website (http://tuvalu.santafe.edu/~aaronc/wsbm/). We varied the number of communities from *k* = 2,…, 8 and repeated the optimization procedure 25 times, each time initializing the algorithm with a different set of parameters.

### Module interaction motifs

A fitted WSBM assigns brain areas to communities. We can characterize the interactions between communities based on their pairwise community densities. The interaction between two communities, *r* and *s*, can be characterized by the community densities:

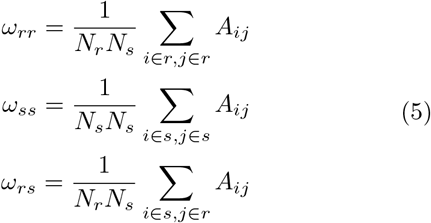

where *N*_*r*_ and *N*_*s*_ are the number of nodes assigned to communities *r* and *s*, respectively. Given these densities, we classify the community interactions as one of three types:

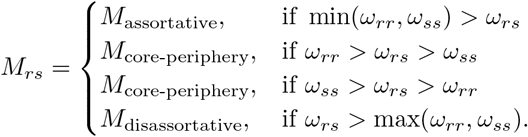

Note that singleton communities have undefined internal densities. Consequently, we ignore singleton communities, treating those brain areas as if they participated in no motifs.

### Mapping interaction motifs to brain areas

Interaction motifs are defined at the level of communities. To map motifs to brain areas we adopt the following procedure. Given a *k*-community partition, each community *r* participates in *k* – 1 interactions, which may represent one or more motif type. Let *M*_*r*_ = {*m*_1_,…,*m*_*k*−1_} where *m*_*s*_ ∈ {“Assortative”, “Disassortative”, “Core”, *and* “Periphery”} denote the set of interaction types in which community *r* participates. Given *M*_*r*_, we can calculate *P*_*rt*_ as the fraction of all *k* − 1 interactions that are of type *t*. These values can be propagated to individual brain regions by letting *P*_*it*_ = *P*_*rt*_ for all *i* ∈ *r*. In practice, repeated runs of the WSBM optimization algorithm can result in different partitions and possibly different interaction motifs. We calculated *P*_*it*_ for each run of the algorithm and averaged the values over all runs to generate for each brain area a single mean value.

### Diversity index

In addition to studying interaction motif classes in isolation, we also characterized the diversity of motif classes in which a brain area participates as the entropy over its interaction motif distribution. In particular, we defined a regional diversity index as 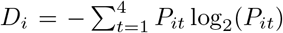. Intuitively, *D*_*i*_ is equal to zero bits if the community to which brain area *i* is assigned participates in only one motif class, and it is equal to 2 bits if *i* participates in all motif classes uniformly.

### Module dominance index

Finally, we defined a community dominance index, which measures for each brain area the motif class that it is most strongly associated with, compared to other brain regions. To calculate community dominance, we first column-normalize *P*_*it*_, so that 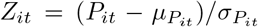. The community dominance is the index, *t* ∈ {1, 2, 3, 4} corresponding to the maximum value of *Z*_*it*_ over all *t*.

### Task state classification

We aimed to classify the task state of subjects based on network measures derived from their community structure. One of the most common classifiers is the decision tree, which partitions a feature space into sub-spaces where each sub-space delineates a discrete class [75]. In our case, the feature space is defined by network measures, and the discrete class is the task state. While decision trees are common and generally useful, single decision trees may be prone to overfitting. One strategy for mitigating this issue is to combine multiple decision trees trained on randomly selected but smaller sub-sets of features [76]. Here, we use this “random forest” approach (implemented in MATLAB as its `treebagger.m` function [77]) to classify subjects’ task states.

We used random forests in two separate contexts. In the first case we defined a series of feature sets at the areal level. That is, each brain area is associated with a five-dimensional feature set comprised of its assortativity, disassortative, core, and periphery expression levels in addition to its community diversity score. For each brain area, we fit a random forest model using these feature sets aggregated across subjects to classify subjects’ task state labels. In our implementation, the random forests comprised 100 decision trees trained on 90% of the data with the held-out 10% serving as test data. Importantly, all train-test splits were performed at the level of subjects, ensuring that the test set included no data from subjects in the training set.

In the second case, we used precisely the same implementation but with feature sets defined based on principal components derived from brain-wide patterns of assortativity, disassortative, core, and periphery expression levels as well as community diversity scores. In this case, the dimensionality of the feature set was user-determined and corresponded to the number of components retained, *P*. We explore how varying P influences our results in Fig. S4.

## AUTHOR CONTRIBUTIONS

This study was designed, carried out, and written by RFB. MAB processed and provided MRI and behavioral data. All authors contributed to the direction of the research and edited the paper.

## ACKNOWLEDGEMENTS

RFB, MAB, and DSB acknowledge support from the John D. and Catherine T. MacArthur Foundation, the Alfred P. Sloan Foundation, the ISI Foundation, the Paul Allen Foundation, the Army Research Laboratory (W911NF-10-2-0022), the Army Research Office (Bassett-W911NF-14-1-0679, Grafton-W911NF-16-1-0474, DCIST-W911NF-17-2-0181), the Office of Naval Research, the National Institute of Mental Health (2-R01-DC-009209-11, R01 MH112847, R01-MH107235, R21-M MH-106799), the National Institute of Child Health and Human Development (1R01HD086888-01), National Institute of Neurological Disorders and Stroke (R01 NS099348), and the National Science Foundation (BCS-1441502, BCS-1430087, NSF PHY-1554488 and BCS-1631550). The content is solely the responsibility of the authors and does not necessarily represent the official views of any of the funding agencies.

Data were provided [in part] by the Human Connectome Project, WU-Minn Consortium (Principal Investigators: David Van Essen and Kamil Ugurbil; 1U54MH091657) funded by the 16 NIH Institutes and Centers that support the NIH Blueprint for Neuroscience Research; and by the McDonnell Center for Systems Neuroscience at Washington University.

---

**FIG. S1.**
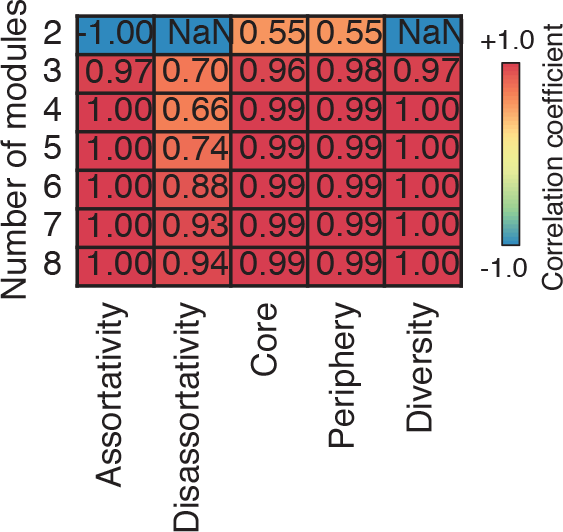
Comparing network measures computed using the first resting state scan (REST1 data) with those computed using the second resting state scan (REST2 data) Columns correspond to network measures and rows correspond to the number of detected communities, *k*. Each cell is the Pearson correlation of REST1 with REST2 computed for a given network measure at a given *k*.

**FIG. S2.**
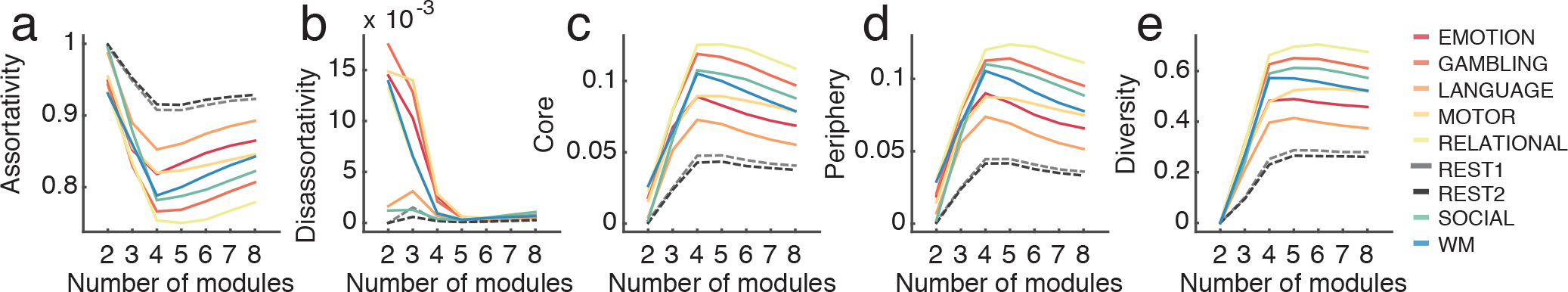
Variation of assortativity, disassortativity, core, periphery, diversity, and node-level true positive rates as a function of *k*. (*a*) Mean assortativity, (*b*) disassortativity, (*c*) core, (*d*) periphery, and (*e*) diversity for each task type as a function of the number of communities, *k*. Note that the assortativity of REST1 and REST2 are consistently greater than that of any of the cognitive tasks. Conversely, core, periphery, and diversity of REST1 and REST2 are consistently smaller than that of any of the cognitive tasks. The colored lines represent the eight scan types and the colored lines while the black lines represent the mean across all tasks.

**FIG. S3.**
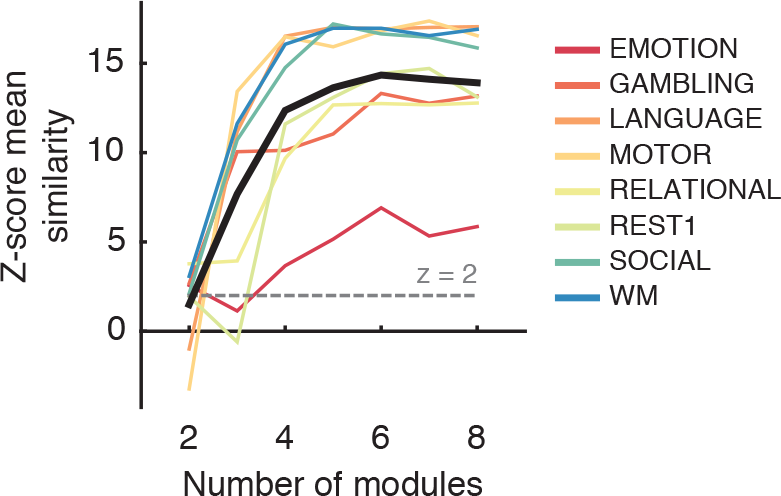
Similarity of node-level true positive rates as a function of *k*. For each *k* and for each task, we generate *N* × 1 vectors of true positive rates. For each task, we calculated the mean similarity of the *k′*th vector with all other *k* = 2,…, 8, where we measure similarity as a Pearson correlation. We compare these empirical mean similarity scores with null similarity scores, generated by randomly and uniformly permuting the elements of the *N* × 1 vector independently (1000 repetitions). The empirical similarity scores are standardized as the *z*-score with respect to this null distribution. Here, we plot *z*-scores for each task as a function of *k*. The colored lines represent the eight scan types while the black lines represent the mean across all tasks.

**FIG. S4.**
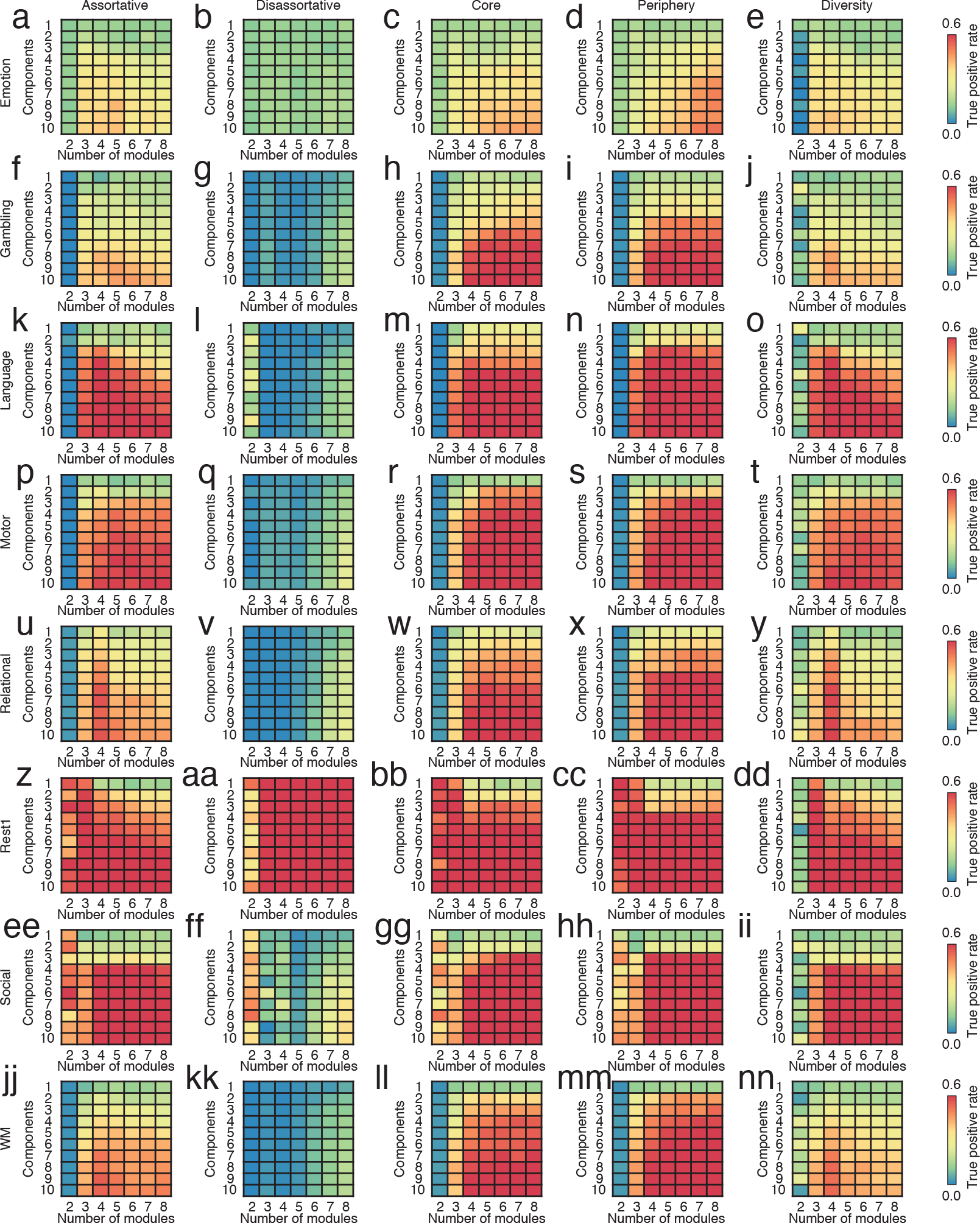
Variation of true positive rate with the number of communities, *k*, and with the number of principal components, *P*, for all network measures and all tasks. Each 10 × 7 matrix shows variation in true positive rate with the number of detected communities, *k*, and the number of principal components, *P*, used for classification. Note that *P* is equivalent to the number of features. In general and intuitively, we find that increasing *P* leads to better performance. As in the main text, we also find that disassortativity performs poorly and that core and/or periphery expression levels outperform other features.

**FIG. S5.**
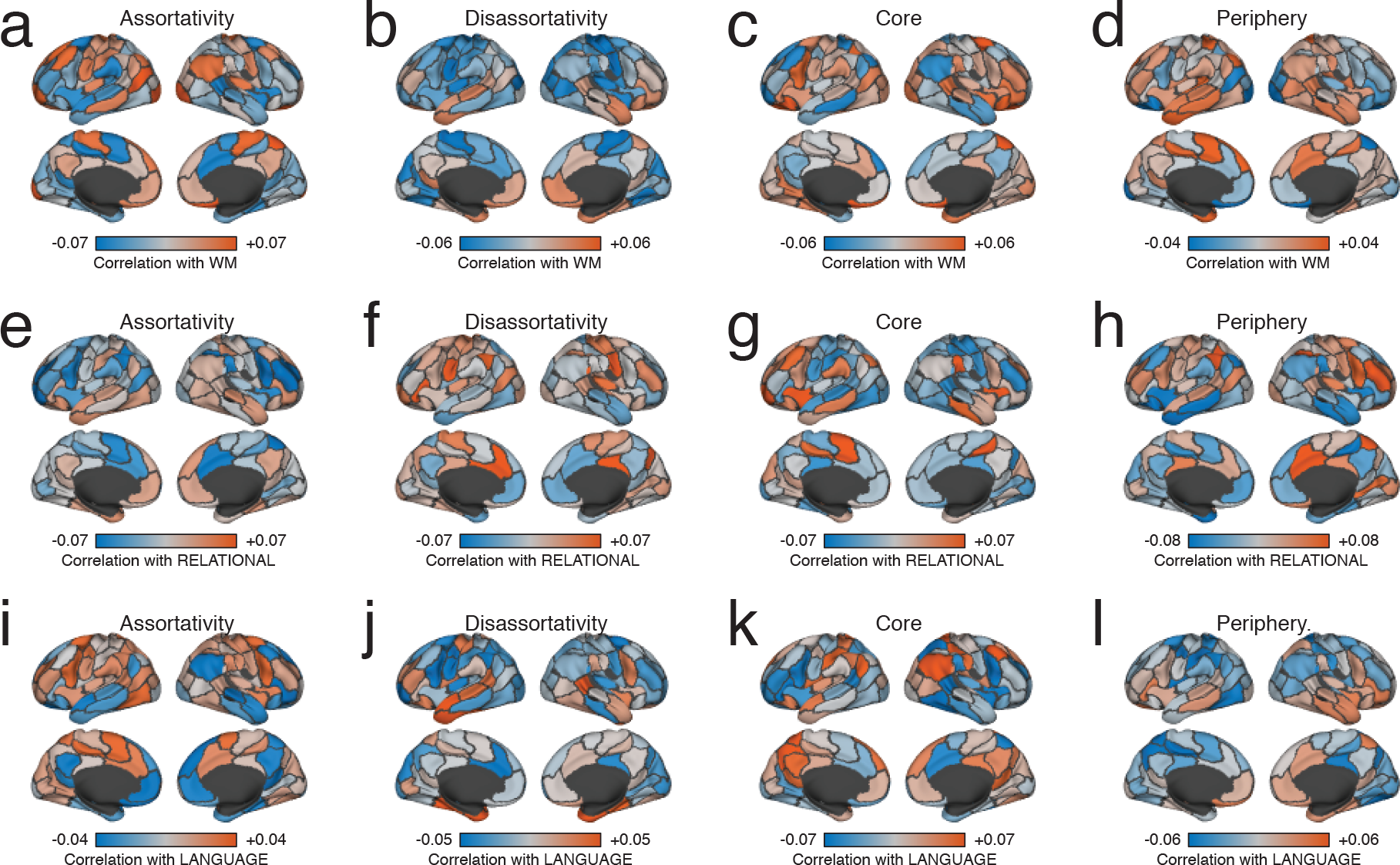
Correlations of network and behavioral measures. In the main text, we plotted the correlation of behavioral measures (SOCIAL) and network measures (computed using REST1) onto the cortical surface. Here, we show similar correlations but for the (*a*-*d*) WM (working memory), (*e*-*h*) RELATIONAL, and (*i*-*l*) LANGUAGE tasks.

**FIG. S6.**
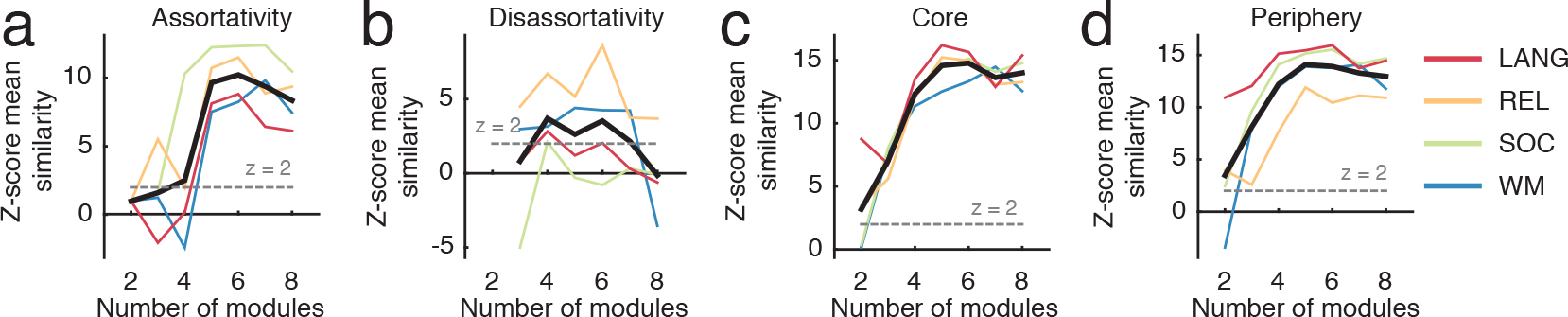
Similarity of brain and behavioral correlations as a function of the number of detected communities. We computed the correlation across subjects of node-level network measures with behavioral measures, resulting in an *N* × 1 vector of correlation coefficients. In the main text, we show correlation coefficients for *k* = 6. Here, and for each network measure, we compute the mean similarity of the *k*′th vector with all other *k* = 2,…, 8, where we measure similarity as a Pearson correlation. We compare these empirical mean similarity scores with null similarity scores, generated by randomly and uniformly permuting the elements of the *N* × 1 vector independently (1000 repetitions). The empirical similarity scores are standardized as the *z*-score with respect to this null distribution. Here, we plot *z*-scores for each task as a function of *k*. In each plot the four colored lines represent the four behavioral measures while the black lines represent the mean across all measures.

